# Benchmarking available bacterial promoter prediction tools: potentialities and limitations

**DOI:** 10.1101/2020.05.05.079335

**Authors:** Murilo Henrique Anzolini Cassiano, Rafael Silva-Rocha

**Author notes:** Correspondence to: Rafael Silva-Rocha, Faculdade de Medicina de Ribeirão Preto, Universidade de São, Paulo, Av. Bandeirantes, 3.900. CEP: 14049-900., Ribeirão Preto, São Paulo, Brazil.

## Abstract

**Background:** The promoter region is a key element required for the production of RNA in bacteria. While new high-throughput technology allows massive mapping of promoter elements, we still mainly relay on bioinformatic tools to predict such elements in bacterial genomes. Additionally, despite many different prediction tools have become popular to identify bacterial promoters, there is no systematic comparison of such tools.

**Results:** Here, we performed a systematic comparison between several widely used promoter prediction tools (BPROM, bTSSfinder, BacPP, CNNProm, IBBP, Virtual Footprint, IPro70-FMWin, 70ProPred, iPromoter-2L and MULTiPly) using well-defined sequence data sets and standardized metrics to determine how well those tools performed related to each other. For this, we used datasets of experimentally validated promoters from *Escherichia coli* and a control dataset composed by randomly generated sequences with similar nucleotide distributions. We compared the performance of the tools using metrics such as specificity, sensibility, accuracy and Matthews Correlation Coefficient (MCC). We show that the widely used BPROM presented the worse performance among compared tools, while four tools (CNNProm, IPro70-FMWin, 70ProPreda and iPromoter-2L) offered high predictive power. From these, iPro70-FMWin exhibited the best results for most of the metrics used.

**Conclusions:** Therefore, we exploit here some potentials and limitations of available tools and hope future works can be built upon our effort to systematically characterize such quite useful class of bioinformatics tools.

## Introduction

Promoter regions are intrinsic DNA elements located upstream of genes and required for its transcription by the RNA polymerase (RNAP) [1]. Thus, the correct mapping of promoters is a critical step when studying gene expression dynamics in bacteria. While the definition of promoters could vary widely, we will consider here as the core elements recognized by the sigma subunit of the RNAP. In *Escherichia coli*, seven alternative sigma factors are responsible for gene expression, while simga70 is the most important one as it is required for the expression of housekeeping genes [2,3]. Therefore, this sigma factor recognizes a consensus region about 35 bp in length with two key elements, the -10 box (with consensus motif TATAAT) and the -35 box (TTGACA) which are separated by 17 ± 2 bp [1,2]. In addition to the core promoter region, other *cis*-regulatory elements can play relevant roles in the regulation of gene expression [4]. In this sense, the production of RNA at the transcription start site (TSS) is the result of the interplay between the core promoter region and the *cis*-regulatory elements [5]. Mapping of functional promoter elements have been performed mostly using low-throughput techniques (such as promoter probing, primer extension, DNA footprinting, etc.) or more recently, by high-throughput experimental approaches (RNA-seq, Genome SELEX, Sort-seq, etc.) [6–11]. However, the rapidly growing number of fully sequenced bacterial genomes greatly exceeds our ability to map promoter elements experimentally. Therefore, diverse computational tools have been created to predict promoters/TSSs at specific genes or at genomic levels.

Some of the first approaches to map promoters have been based on the use of position-weighted matrices of -10 and -35 box motifs, taking into account the distribution of the spacer length between them and their distance from TSS [12,13]. Yet, over the past years, a growing number of computational strategies have evolved in complexity. Notable novel approaches raised, such as sequence alignment-base kernel for support vector machine [14,15], profiles of hidden Markov models combined with artificial neural networks [16] or weighted rules extracted from neural network models [17]. Also, new ways to extract information from DNA sequences to perform predictions have appeared. Thus, there are now several numerical representations of DNA sequences which each one carry its own properties [18–20], such as methods that uses k-mer frequencies or variations [21,22] and other methods that includes physicochemical properties of DNA [23].

Recently, machine learning (ML) techniques have been used to obtain insight from different sources from diverse biology fields (an extensive survey can be seen in Libbrecht and Noble, 2015; Camacho *et al.*, 2018; Zou *et al.*, 2019), and in the past few years this has been applied to the recognition of promoters, TSSs and regulatory sequences. Among most of the ML algorithms used for this purpose we can mention Support Vector Machine [27], Neural Networks [28], Logistic Regression [29], Decision Trees [30], and Hidden Markov models [31,32]. Despite the existence of all these modern techniques, promoters cannot always be inferred based on their sequence only, and, currently, we have no clue on how efficient these tools indeed are. This occurs since each new tool is validated without the use of standardized datasets or methods, making difficult to compared novel emerging alternatives with the current state of art. In this work, we summarize general aspects of the available promoter prediction tools, exposing in a comparative manner their main strong and weak features. For this, we compared the performance of these tools using experimentally-validated promoters from *E. coli*. Unexpectedly, we show that some very popular tools such as BPROM performed very poorly compared to tools created over the last two years. We hope our results can help both community users to choose a suitable tool for their specific applications, as well as developers to construct novel tools overcoming key limitations reported here.

## Results and discussion

### Describing the tools: methods, availability and usability

In this section, we present a succinct explanation of each methodology (**Table 1**) as well as the usability information about their use requirements, acceptable file types, etc. (**Table 2**). Below, we described briefly for each tool how they have been built and some of the main features.

**Table 1.**
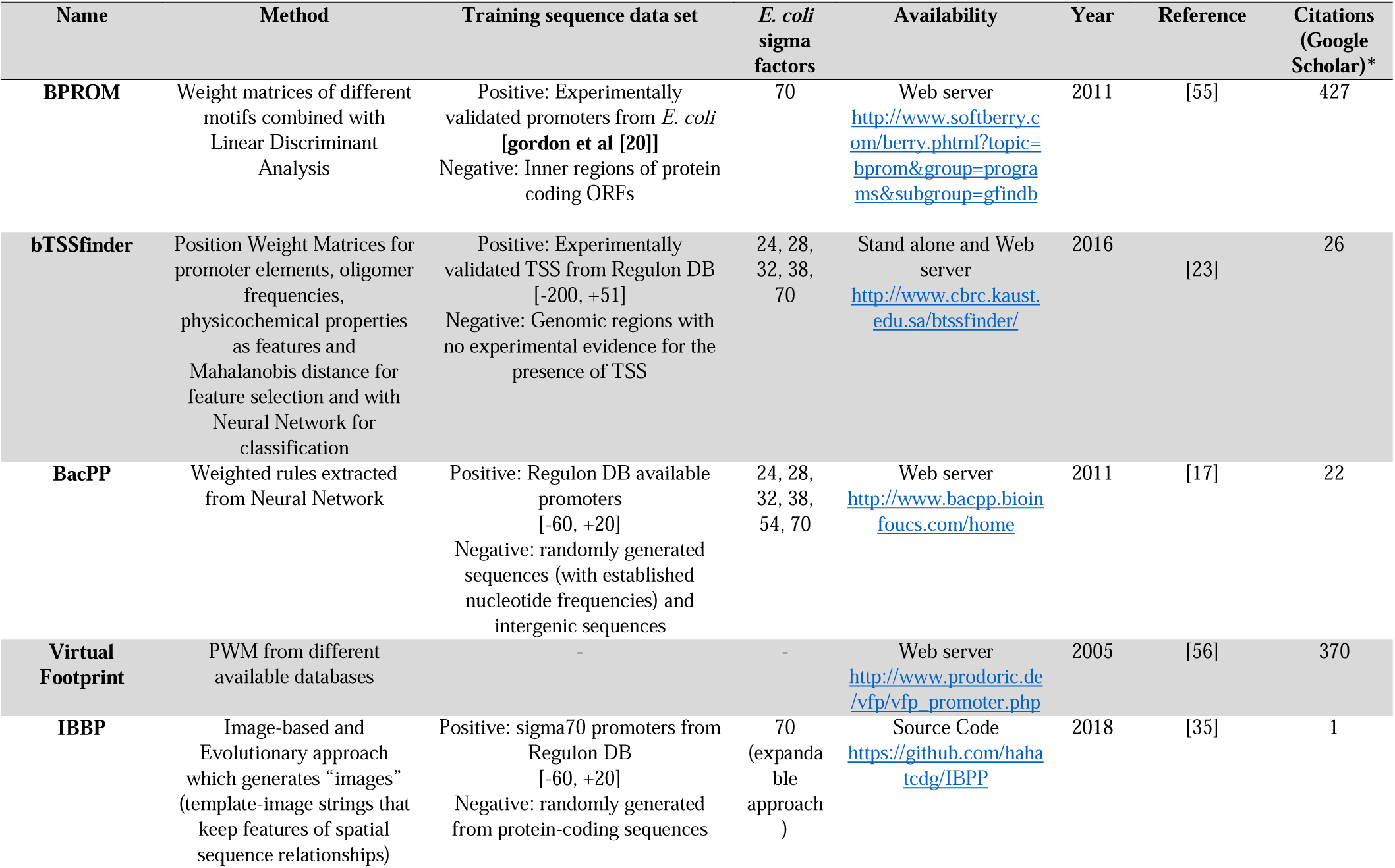
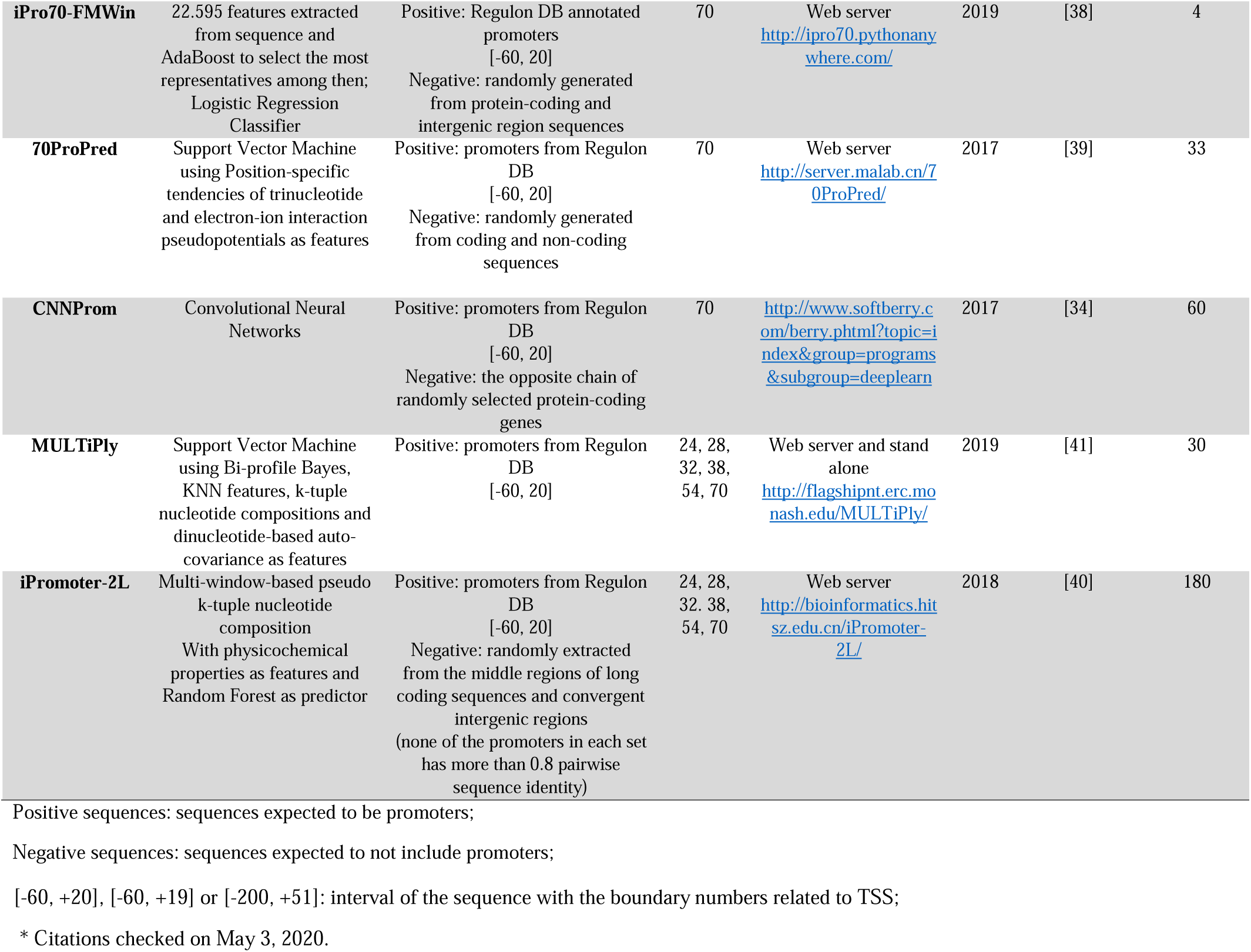
General information of the tools used here.

**Table 2.**
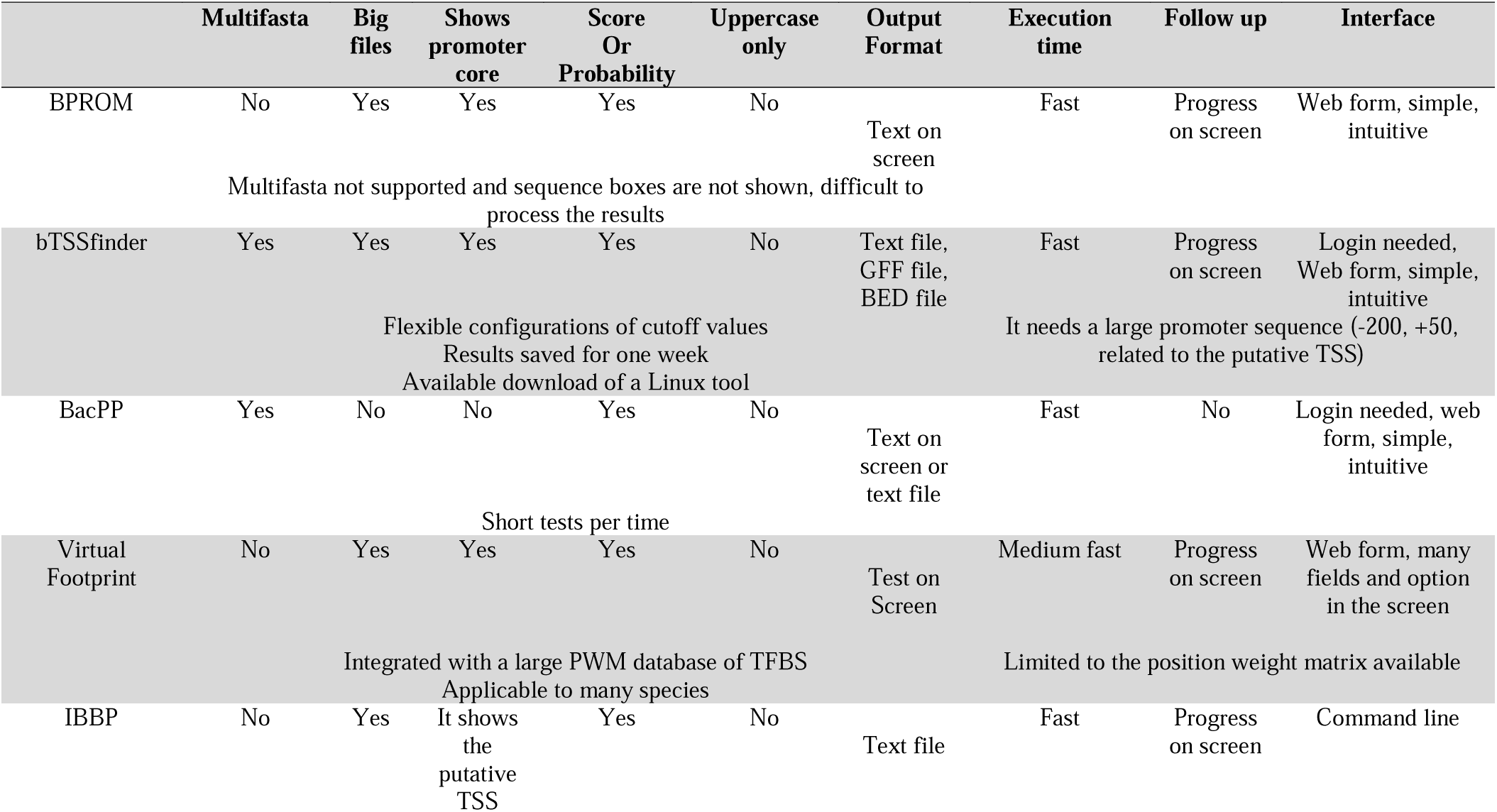
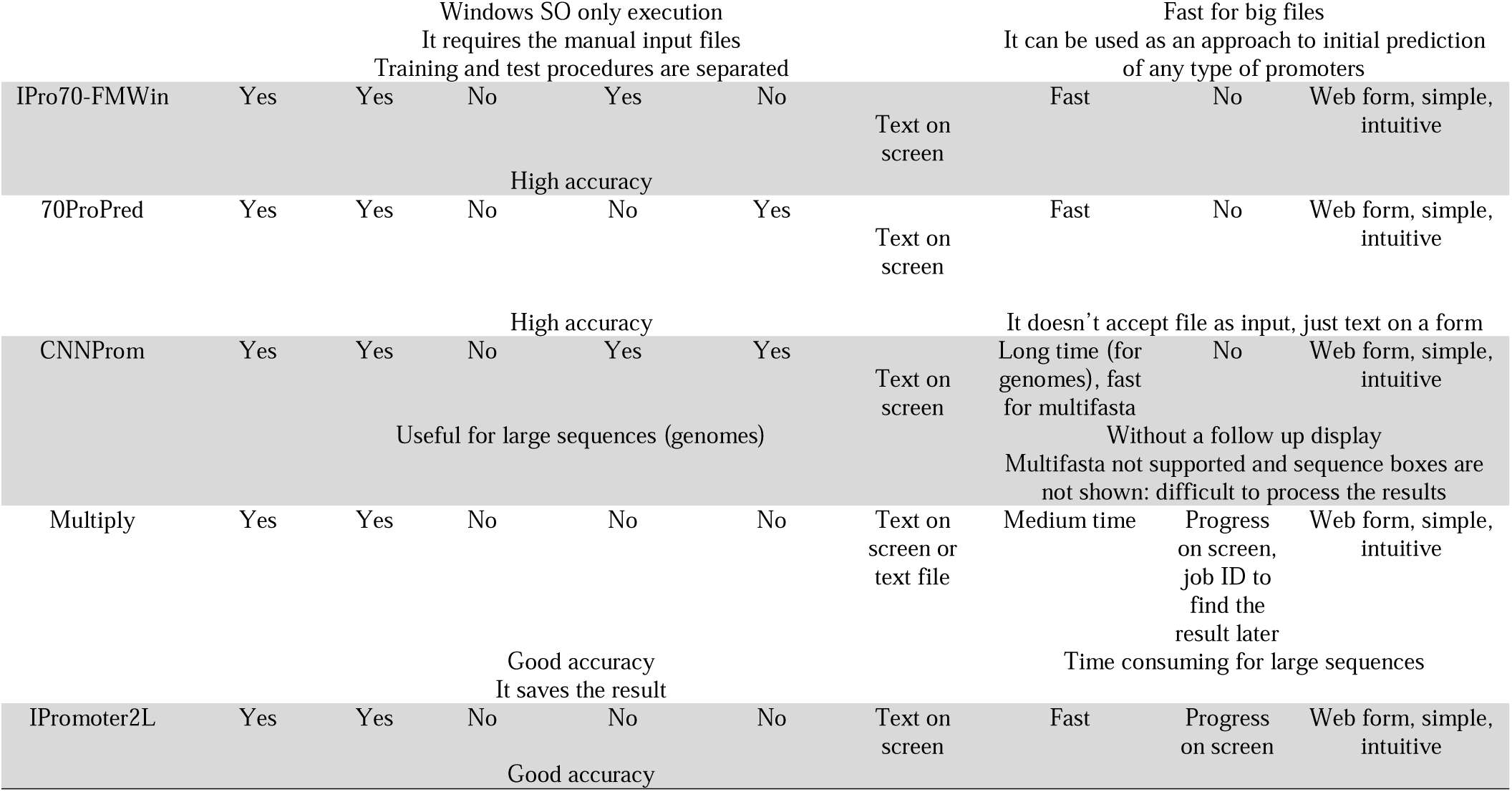
Usage characteristics of the tools analyzed here.

*BPROM* [33] was developed as a module of an annotation pipeline for microbial sequences in order to find promoters in upstream regions of predicted ORFs. To train the model the authors used a dataset of experimentally validated promoters from elsewhere [14]. They applied linear discriminant analysis to discriminate between those promoters and inner regions of protein-coding sequences. As attributes they used 5 position weighted matrices of promoter conserved motifs and they also consider the distance between the –10 and –35 boxes and the ratio of densities of octa-nucleotides overrepresented in known bacterial transcription factor binding site (TFBS) relative to their occurrence in coding regions. This tool is available as a web application and users can submit a local file or past the sequence in the web form. It quickly returns the results in the screen with the possible –10 and –35 boxes of predicted promoters and its position in the submitted sequence.

*bTSSfinder* [23] is a tool that predicts putative promoters for different sigma factors in *E. coli* and Cyanobacteria. Its positive dataset consists of experimentally validated *E. coli* TSS from Regulon DB and different experimentally mapped Cyanobacteria TSS provided by several works. Its negative dataset consists of genomic regions where there is no experimental evidence for the presence of TSS. They started with 30 features distributed between these types: promoter element motifs (PWMs), distance between the elements, oligomer scores, TFBS density and physicochemical properties. The final set of features were selected by evaluating the predictive power of these features by calculating Mahalanobis distance and used to training a neural network. This tool is available as a web application or as a stand-alone tool for Linux. In the website, an email is needed to login and the results are saved for a week.

*BacPP* [17] is a prediction tool to find *E. coli* and other Enterobacteriaceae promoters. As positive dataset the authors used promoter sequences from Regulon DB for 6 different sigma factors in *E. coli* and other Enterobacteriaceae promoter sequences obtained from several works. In its negative dataset they used 2 approaches: a) random sequences generated with a probability of 28% for nucleotides adenine and thymine and 22% for cytosine and guanine; b) random selected intergenic regions. Each nucleotide of these sequences was transformed in binary digits and used to train neural networks. To use this tool, the user must create a login in the website then past the sequences or fasta file according their model and select the sigma factors of interest.

*CNNProm* [34] is a web tool that can predict prokaryotic and eukaryotic promoters from big genomic sequences or multifasta files. In the case of *E. coli* promoters, authors took the sequences from Regulon DB and the negative control (non-promoter sequences) were randomly selected from the opposite chain of coding regions in genomes. Each of these sequences were transformed in a binary four-dimensional vector and used directly as features to train a convolutional neural network. To use this predictor, users must enter the sequences or the file in the website and choose the organism model.

*IBBP* [35] is a stand-alone application that implements a new approached called “image-based promoter prediction”. This approach consists in generating multiple “images”: template strings carrying possible features/elements presented in promoters and their spatial relationships. The image generation and selection are made by applying an evolutionary approach and calculating the similarity of these images in a set of *E. coli* sigma70 promoters. The authors measured the accuracy of the tool by analyzing the set of promoters and protein-coding sequences. To use this software, it is necessary to download the executable files, execute the evolutionary algorithm with the promoters of interest and then implement the classifier software, which uses the result model generated in the previous step.

*Virtual Footprint* [36] is a web framework for prokaryotic regulon prediction. This framework makes use of several PWM provided by PRODORIC [37] and other PWM from other sources. To make the prediction, it is necessary to upload a DNA sequence or a fasta file, select different PWM for core promoter elements or other transcription factor binding sites and set some parameters.

*IPro70-FMWin* [38] is a web application for sigma 70 promoter prediction. Its training dataset consists of sigma 70 promoter sequences from dataset Regulon DB 9.0 and sequences randomly chosen from coding and intergenic regions of *E. coli*, as positive and negative datasets, respectively. For feature extraction, 22,595 sequence-based features were generated for “multiple windows”, i.e., different regions of the promoter sequence. These features include, for example, different kinds of k-mer and g-gapped k-mer compositions, statistical and nucleotide frequency measures. Among the machine learning methods tested be the authors, logistic regression achieved better results. They also applied AdaBoost technique for feature selection in order to improve prediction.

*70ProPred* [39] was built using sigma 70 promoter sequences from Regulon DB 9.0 and randomly generated sequences from coding and non-coding regions of *E. coli* genome to train a support-vector machine (SVM) model. The attributes generated from the sequences were position-specific trinucleotide propensity and electron-ion interaction pseudopotentials of nucleotides, considering single or double stranded DNA, to reveal trinucleotide distribution differences between the samples and represent interaction of trinucleotides, respectively.

*iPromoter-2L* [40] is an online tool that provides prediction for all *E. coli* sigma promoters. This method has two “layers” of classification applying random forests, first it resolves whether a given sequence is a promoter and then it selects the sigma factor class. For model training, the authors used experimentally confirmed promoter sequences from Regulon DB 9.3, as positive dataset, and randomly extracted sequences from the middle regions of long coding sequences and convergent intergenic regions, as negative dataset. It is important to emphasize that sequences with more than 0.8 pairwise sequence identity, for a given sigma factor promoter dataset, were removed in order to reduce identity biases. Their feature extraction was based on multi-window-based pseudo K-tuple nucleotide composition, which consists in a sliding window, extracting and encoding physicochemical attributes of different regions of a given sequence.

*MULTiPly* [41] web application provides promoter prediction for all *E. coli* sigmas. To train their model, the authors used experimentally validated promoter sequences from Regulon DB for all type of sigma factor in *E. coli*. Their feature extraction was divided in two types, the first one was used to represent global features, applying bi-profile bayes and KNN features, and the second-one to represent local features, applying k-tuple nucleotide composition (sequence-based feature) and dinucleotide-based auto-covariance (which considers physicochemical properties). This method also performs two steps of classification: first it resolves whether a given sequence is a promoter or not, then it decides which class of sigma promoter it belongs. The authors used SVM method for classification and F-score method for feature selection.

These last 4 web tools (*IPro70-FMWin, 70ProPred, iPromoter-2L* and *MULTiPly*) have a similar way to use, accepting multifasta formatted sequences on a simple web form and returning the results on the screen. Particularities and a summarization of the approaches discussed above are present on **Table 1** and **2**.

### Analyzing the performance of promoter prediction tools

In order to compare the performance of the promoter prediction tools presented above, we analyzed the positive and negative datasets as described in the methods section. From the ten algorithms selected, BacPP could not be tested with our entire dataset, because multifasta files were not supported, and Virtual FootPrint produces a large number of predicted –10 box for sigma 70 in both positive and negative datasets, a number that greatly exceeds the number of sequences analyzed. Thus, these two tools were not considered in further analyses. Among the remining 8 algorithms, 5 achieved more than 50% of correct classification on positive dataset, while 6 correctly classified 50% of the negative dataset (**Figure 1A**). The best performance was observed for CNNProm (94.8% TP), followed by iPromoter-2L (83.8% TP), 70ProPred (89.7% TP), iPro70-FMWin (94.5% TP) and MulTiPly (81.2% TP). When we compared the performance parameters (accuracy, MCC, sensitivity and specificity), we observed that four of them (CNNProm, iPromoter-2L, 70ProPred and iPro70-FMWin) presented the best performance, while MulTiPly only scores high for sensitivity (**Figure 1B**). Therefore, we can observe MCC values close to zero for the remaining 4 tools (<u>MulTiPly</u>, bTSSFinder, BPROM and IBPP), indicating that these tools performed close to random classifications. It is interesting to notice that BPROM, a widely cited and used tool, presented the worst results together with bTSSFinder and the IBBP, but also presented the fewer FPs. We also found that IBBP’s method based in evolutionary approach classified random sequences as promoters more than in real promoter dataset (i.e., it displays higher FP rate than TP). From the analysis presented in **Figure 1**, we can observe that iPro70-FMWin performed best due to a small number of FP and the overall bests results of all metrics used (**Figure 1B**).

**Figure 1.**
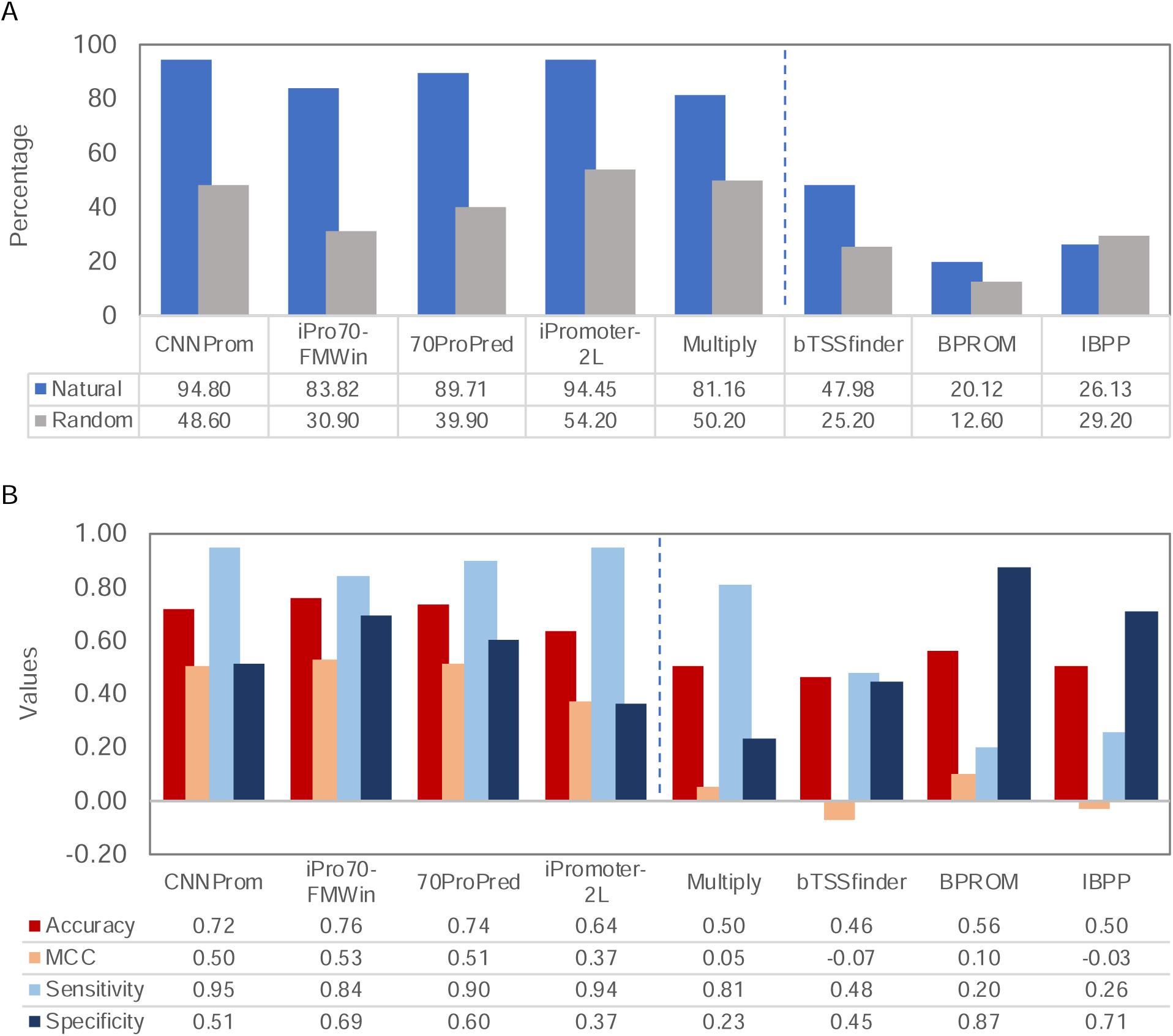
Analysis of the performance of promoter prediction tools. **A**) Percentage of sequences predicted as sigma 70-dependent promoters in both datasets. The percentage of correct classifications of experimental promoters (blue) and percentage of misclassified random sequences (gray) are presented. Vertical bashed line separates the five best to the three worse tools analyzed. **B**) Metrics used to evaluate the performance of the tools. Note that MCC values range from -1 to 1. It is important to emphasize that two tools presented highest sensitivity associated with a low specificity, i.e., tools usually perform good classifications for real promoters and high misclassification of random sequences. Vertical dashed line divides the four best from the four worse tools.

We next performed a hierarchical clustering analysis using the results from the five tools that presented the best results. As can be seen in **Figure 2A** for the positive dataset, results obtained with iPromoter-2L were more correlated with CNNProm outcomes, since both produced the larger set of TP predictions, while iPro70-FMWin was more related to 70ProPred. In general, 573 sequences (62.2%) where correctly classified by all 5 algorithms (**Figure 2B**). When we analyzed the negative dataset (constructed with random sequences), we do not observe a clear clustering since each tool presented a different level of FP, with the lowest level observed for iPro70-FMWin (**Figure 3A**). In this case, only 102 sequences (10.2%) were incorrectly classified as promoter by all tools, indicating that each algorithm has specific features to equivocally classify the random sequences. It is worth mentioning that the best three tools (CNNProm, iPro70-FMWin and 70ProPred) are from 2017 to 2019, indicating that, as expected, promoter prediction algorithms are evolving through the years. Taken together, these results indicate that 4 out of 8 tools analyzed here display equivalent predicting power to identify true promoter sequences, while widely used tool BPROM exhibit a reduced predictive capability.

**Figure 2.**
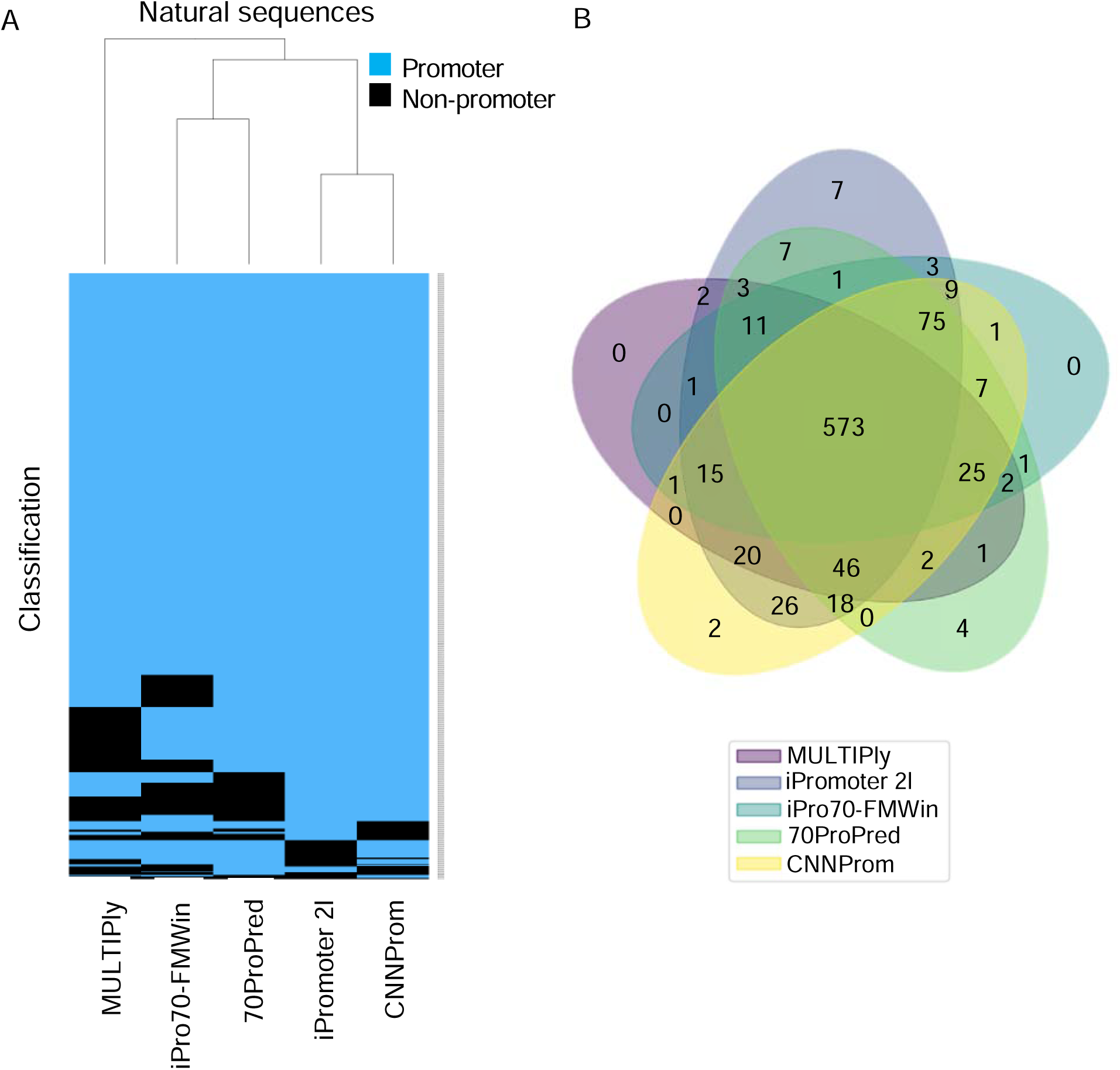
Analysis of tool performance in positive dataset (natural sequences). **A**) Hierarchical clustering of DNA sequences classified as promoters (blue) or non-promoters (black). **B**) Venn diagram representing the number of sequences predicted as promoters from panel A.

**Figure 3.**
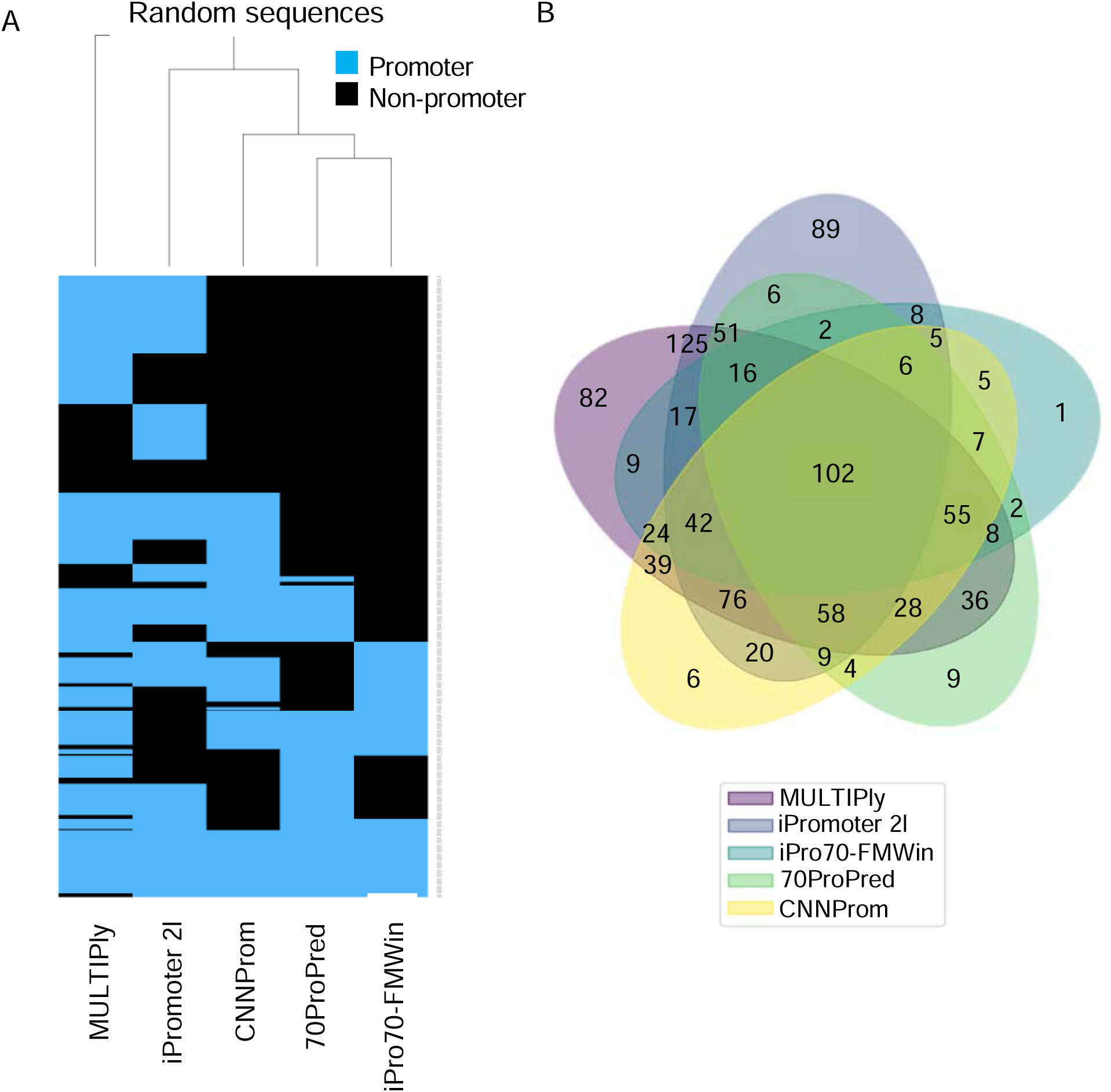
Analysis of tool performance on negative dataset (random sequences). Hierarchical clustering of DNA sequences classified as promoters (blue) or non-promoters (black). **B**) Venn diagram representing the number of sequences predicted as promoters from panel A.

### Identification of promoter features identified during the analyses

As presented above, we observed a high degree of similarity between the best tools for the identification of true promoters, but a lower overlap on random sequences equivocally classified as promoters. This could indicate that each algorithm might identify different features to assign a sequence as promoter. In order to further investigate this process, we analyze the information content from the sequences identified as promoters from the positive and negative dataset for the top 5 tools analyses here. The results of these analyses are presented as sequence Logo on **Figure 4** and **5** for positive and negative dataset, respectively. As can be seen on **Figure 4**, TP sequences identified by all 5 algorithms display the same consensus sequence that resembles a strong canonical -10 box from sigma70 promoters [2]. It is worth noticing that the information content was higher for iPro-70-FMWin (up to 0.4 bits), which also displayed the best performance according to the metrics used here. However, when we analyzed the data from promoters identified in the random sequences, we could see a much fuzzy signal for MulTiPly, iPromoter-2L and CNNProm, which were the three tools with the highest FP rate from the top five tools (**Figure 5**), indicating that the rich A (adenine) and T (thymine) frequencies play a role on false positive classification. We also could observe that, in the case of random sequences predicted as promoters for all algorithms, we obtained a more evident -10 box motif and it still shows high A and T influences (**Figure S1**). This implies that these tools are sensitive to AT content, which makes sense since iPromoter-2L and CNNProm were trained on coding sequences as negative controls [34,40]. On the other hand, 70ProPred and iPro-70FMWin, which presented the lower FP rate, presented a clearer - 10-like signals similar to those identified on the positive sequences, although with lower information content. This could reveal that these tools are classifying sequences which resemble true promoters, and we could not rule out the possibility that some of these random sequences could in fact display promoter activity in *E. coli* if tested experimentally. Taken together, these results indicate that high rates of FP observed for some of these algorithms could be due to the use of unrealistic control sequences (such as coding regions) that could make the algorithms sensitive to AT-rich regions, highlighting the importance of choosing appropriated non-promoter sequences to train these tools.

**Figure 4.**
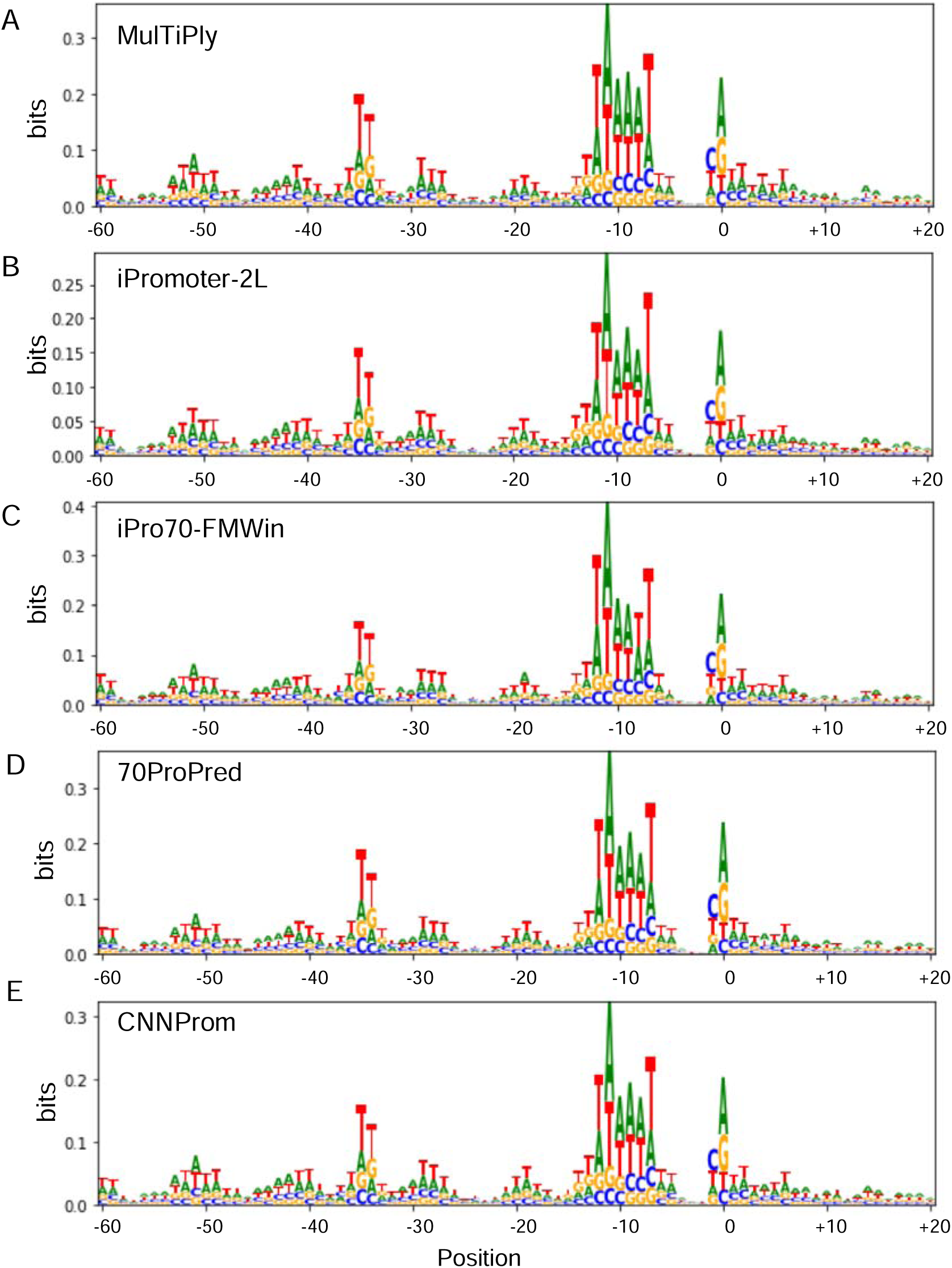
Analysis of information content of DNA sequences identified as promoters on the positive dataset (natural sequences). The sequence logos are shown for sequences predicted as sigma 70-dependent promoters by **A**) MulTiPly, **B**) iPromoter-2L, **C**) iPro70-FMWin, **D**) 70ProPred and **E**) CNNProm.

**Figure 5.**
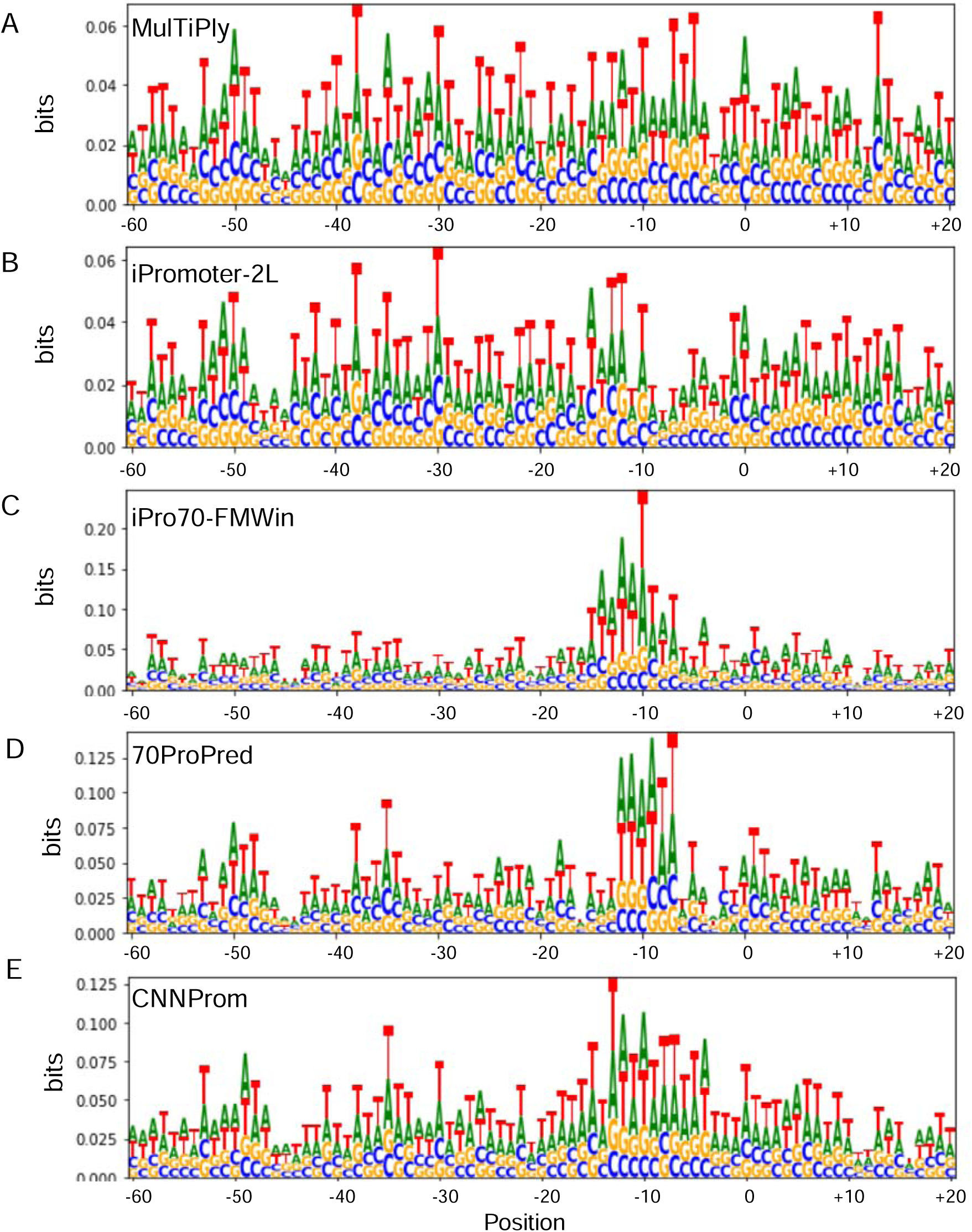
Analysis of information content of sequences identified as promoters on the negative dataset (random sequences). The sequence logos are shown for DNA sequences predicted as sigma 70-dependent promoters by **A**) MulTiPly, **B**) iPromoter-2L, **C**) iPro70-FMWin, **D**) 70ProPred and **E**) CNNProm.

### Conclusions

In this work, we performed a benchmark analysis of promoter prediction tools performance using well-characterized promoter sequence and random sequences. As can be seen from the results above, new tools have emerged with enhanced performance compared to widely used ones. Although best performing tool uses just sequence based features (a result that corroborates with Abbas, Mohie-Eldin and EL-Manzalawy, 2015), in general, algorithms using feature extraction that combines attributes derivate from sequence together with physicochemical properties of DNA achieved better results. It is also clear from our results that choosing the appropriate control (or negative) dataset to construct these algorithms is crucial to avoid false positives. Therefore, coding sequences or sequences with different features renders the tools AT-sensitive, increasing the false positive rate. Furthermore, we still need an experimentally well-validate non-promoter dataset to faithfully use as negative controls in these predictions, but these sequences are not available yet. In this sense, we expect that the growing number of high-throughput experiments could become a great source of data to create novel datasets to train new tools for promoter prediction in the future. Another complication to this subject comes from recent evidence showing that just one mutation in random sequences could lead to constitutive transcription *in vivo*, indicating that transcription is indeed a robust process [43]. Therefore, future attempts have to be made to create complete datasets with very similar promoter/non-promoter sequences in order to train next generation tools.

Additionally, several prior information could be incorporate in prediction methods to improve the final tools. For instance, interrelation between the UP element and a subunit of RNAP were found to play a role on transcription initiation and promoter activity [44] and switch preference of sigma factors in promoters [45]. In addition, specific nucleotide composition and motifs between -10 and -35 boxes leading to different DNA curvatures were found to influence transcription initiation and promoter activity [46,47]. Additionally, more than 300 proteins in *E. coli* are predicted to bind DNA and half of them have their function experimentally characterized [1]. These proteins could thus impact promoter activity *in vivo* and their binding sequence preferences could influence promoter discovery. Finally, recent studies with genomic SELEX show that the number of transcription factor binding sites (TFBS) annotated in databases is underestimated [11]. Thus, it is worth to notice that a promoter is a complex entity that requires a large number of elements, making the transcription observed *in vivo* for a specific DNA element could be due to a number of interplaying factors which perhaps could not be predicted using a single tool (**Figure 6**). One final remark is that the extensive majority of algorithms have been created using datasets of promoters from just one bacterium: *E. coli*. Consequently, since each organism has its particularities in terms of DNA binding proteins and sigma-factor elements, we are still far away from having a prediction tool that can be used for several organisms. In order to accomplish that, we would require extensive promoter datasets from several microorganisms to construct multipurpose prediction tools. Lastly, we hope the approach and metrics used here can contribute to future studies aimed to construct improved promoter prediction tools.

**Figure 6.**
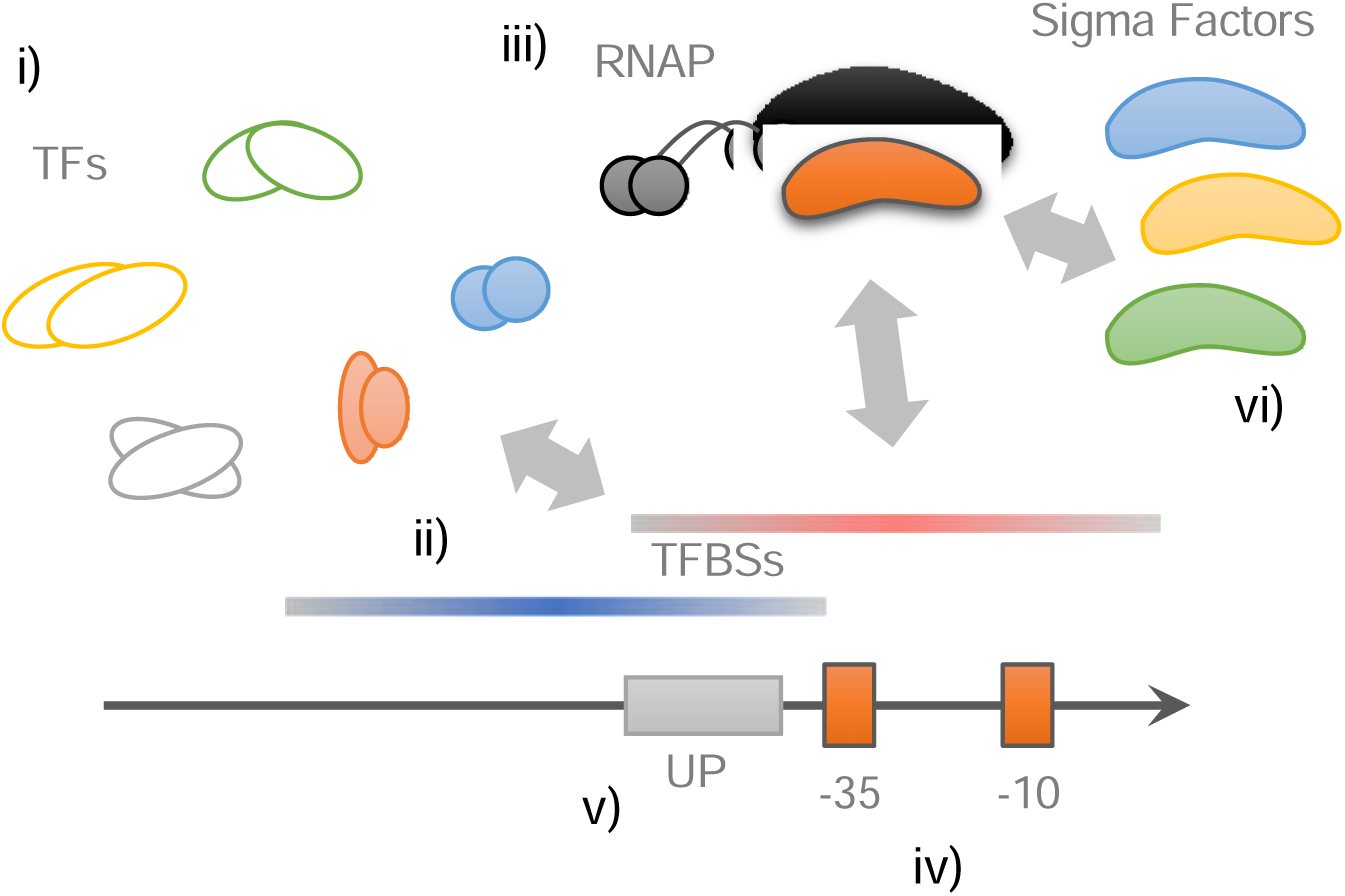
A putative model for a bacterial promoter region, including a range of experimental attributes. **i)** More than 300 proteins (TFs) in *E. coli* are predicted to bind DNA and there is a lack in experimental characterization [1]. **ii)** Recently, high-throughput studies, such genomic SELEX, are showing a large number of possible TFBS on genomes [11], which may impact on the composition of promoter sequences. These regions can have positive (blue regions) or negative (red regions) effect on promoter activity. **iii)** RNAP requires a sigma factor to be recruited to the promoter sequence and each sigma factor possess a preference for a specific motif on DNA [1]. **iv)** Nucleotide composition and motifs between -10 and -35 boxes influences transcription initiation and promoter activity [46,47]. **v)** Interrelation between the UP element and a subunit of RNAP were found to play a role on transcription initiation and promoter activity [44], and the UP element can switch preference of sigma factors in promoters [45]. **vi)** The same promoter sequence can possibly respond to diverse sigma factors, according experimental characterizations and *in silico* approaches [36,57,58].

## Material and methods

### Selecting promoter prediction tools

We started this work by searching in the literature recent and available prediction tools for *E. coli* promoters. For each case, when a tool was available online or by software download, we selected it to posterior analysis. **Table 1** shows the summarized information about the tool methodology (i.e. implementation, approach used, or process performed), the sigma factors it can predict, the available format and the access links. All these descriptions have been extracted from the original papers describing the tools. Next, we analyzed some usability features of the tools (such as the file format accepted as input, maximal allowed file size, the output format of results, etc.) as summarized in **Table 2**. Then, we selected the ones that accepted our complete dataset in multifasta format as input to perform a comparative analysis.

### Promoter datasets used for the analyses

In order to compare each selected tool, we used an experimentally validated promoter dataset for the well-studied *E. coli* K12 (?) which are dependent on sigma70, as available in the curated database Regulon DB 10.5 [48]. We used only sigma70-dependent promoters since they are mostly well-characterized in bacteria, and consequently most tools have been developed to recognize this class of elements. Thus, our so-called positive dataset was formed by 865 natural sequences extracted from Regulon DB and classified as having a strong evidence/confidence level. Additionally, we used a negative promoter set consisting of 1000 randomly generated sequences with a nucleotide distribution similar to that encountered in the 865 natural sequences, which was constructed with an *ad hoc* script written in *Python*. We chose this strategy for two reasons: (i) generating a negative dataset with this approach allows us to assess the tools’ capacity to distinguish real promoters from random sequences, and, (ii) to the best of our knowledge, there are no experimentally validated negative promoter dataset available. Also, it is important to stress that many tools, such as BPROM, 70ProPred and iPro-70FMWin, used coding and intergenic regions as control (negative) sequences, but this is not appropriated since coding and non-coding regions have clearly different nucleotides compositions and structural properties [49,50]. In our datasets, the sequences have 81 bp, since most tools consider and requires as input 60 bp upstream and 20 bases downstream the putative TSS (the region interval [-60, +20]). In the case were the tool required the entire genome, we used *E. coli* K12 MG1655 genome (GenBank U00096.3) and when a tool required a bigger interval than [-60, +20] bp, we extracted the additional sequence from this same genome. The two datasets (natural and random) used here are available as Supplementary data.

### Defining the metrics for promoter analysis

The true promoter (positive) and random (negative) datasets were used to measure the current tools’ capacity to make correct identification of promoter sequences (it is important to emphasize that we are not using our datasets to re-train and test each methodology). Thus, the results were evaluated comparing accuracy and Matthews Correlation Coefficient (MCC, [51]), calculated as the following equations, respectively:

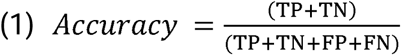

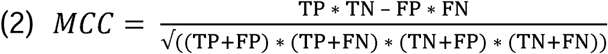

where TP (true positive) are the natural sequences classified as promoters, TN (true negative) are the random sequences classified as non-promoters, FP (false positive) are the random sequences classified as promoters and FN (false negative) are the natural sequences classified as non-promoters. We adopted MCC because it is a metric that deals with unbalanced datasets (i.e. differences in the number of instances in negative and positive datasets), avoiding biases. It achieves high scores only if TP and TN are high, considering both types of correct classification in a single metric, and it has been shown that for this type of binary classification (e.g. promoter/non-promoter) it is more efficient and less over-optimistic [52].

Sensitivity and specificity scores were also used to give a sense of correct classification of promoters and non-promoters, and are defined as follows:

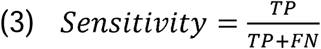

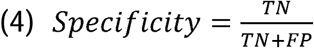

Unlike accuracy, sensitivity and specificity that range from 0 to 1, MCC ranges from –1 (the worst predictor) to 1 (the best predictor) and 0 corresponds to a ‘random’ predictor. By testing the tools with our synthetic random dataset, we can measure whether those tools have overfitting with their test datasets and by testing our positive dataset (with a strong experimental evidence) we are measuring underfitting, once some of our positive sequences probably have already used to train the tools algorithms [24]. As some of the tools also predict promoters for other sigma factors, to be able to classify all predictions as correct or wrong, we considered random sequences classified as any sigma class promoter as FP and a sigma70 sequence classified as any other class of sigma promoter as FN. This does not mean that a sigma70 promoter classified as another sigma factor cannot respond this sigma or even to sigma 70, *in vivo*, as we discuss later.

### Data representation

For data representation, heatmaps were created by using the R package Heatmap.2 [53], with default method and using Jaccard distance method to deal with our binary characteristic vector of 1 (correctly classified) and 0 (wrongly classified) obtained from the tools’ results. The Venn diagrams were made by using the Python library matplotlib-venn (https://pypi.org/project/matplotlib-venn/). The logos of count matrices, probability matrices, position weight matrices and information matrices were constructed by using Logomaker Python library [54]. As every result generated by the tools has different formats, these were pre-processed using a text editor or *ad hoc* Python scripts. All the scripts we used to generate our data and perform the analysis as well as the datasets used are available for download as supplemental files.

## Additional files

Additional file 1: Fasta sequence of natural promoters.

Additional file 2: Fasta sequence of random promoters.

## Ethics approval and consent to participate

Not applicable

## Consent for publication

Not applicable

## Availability of data and materials

The datasets used and/or analysed during the current study are available from the corresponding author on reasonable request.

## Competing interests

The authors declare that they have no competing interests

## Funding

This work was supported by the Sao Paulo Research Foundation (FAPESP, award # 2012/22921-8). MHCA was supported by FAPESP Fellowship (award # 2019/06672-7).

## Authors’ contributions

MHCA and RSR conceived the work. MHCA performed the analysis. MHCA and RSR analyzed the results and written the manuscript. All authors read and approved the final manuscript.

## Acknowledgements

The authors are thanks to lab colleagues for insightful comments on this work, and to Dr. María Eugenia Guazzaroni for comments on the final version of the manuscript.

## Supplementary Figure Legends

**Figure S1.**
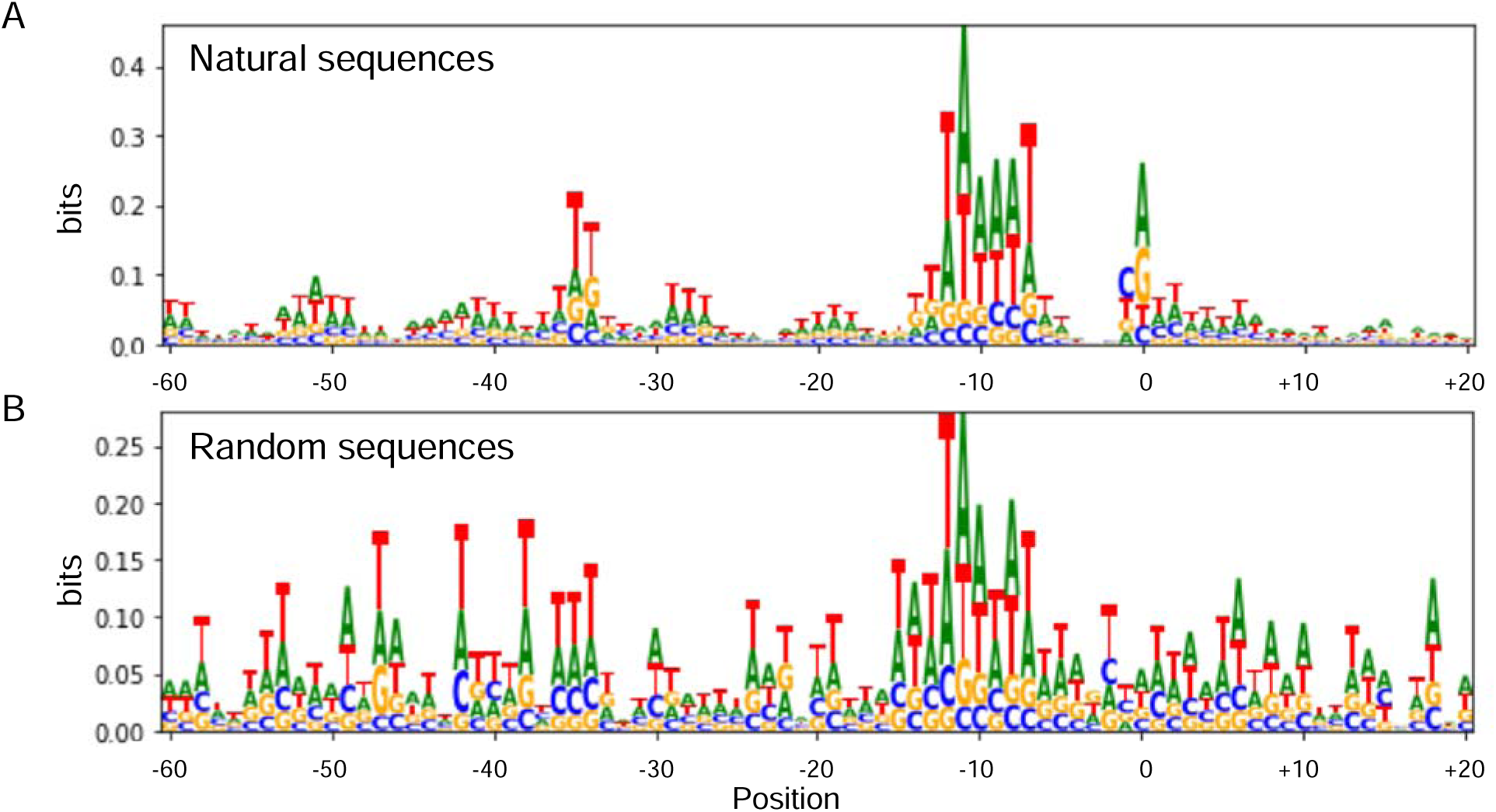
Analysis of information content of sequences predicted as promoters. Sequence logo of the **A)** natural sequences and **B)** random sequences predicted as sigma 70 promoters by all tools. Note that, in the case of natural sequences we observe the same motif pattern that we found in Figure 4, and, in the case of random sequences, the bits range had improved and a -10 box motif become more evident.

## Notes

### Competing Interest Statement

The authors have declared no competing interest.

